## Supplemental Figures for "Zmiz1 is a novel regulator of lymphatic endothelial cell gene expression and function"

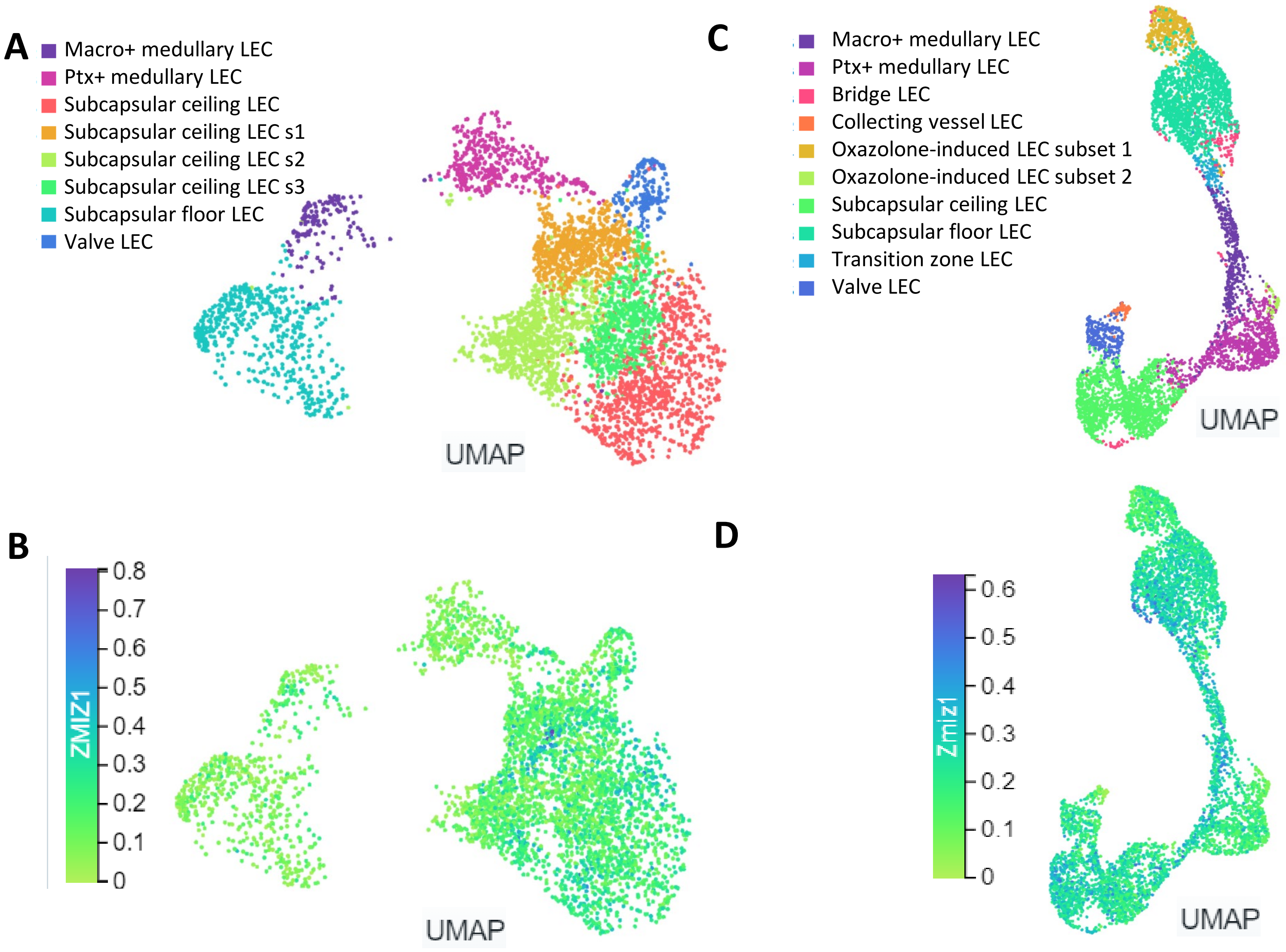
A
Macro+ medullary LEC
Ptx+ medullary LEC
Subcapsular ceiling LEC
Subcapsular ceiling LEC s1
Subcapsular ceiling LEC s2
Subcapsular ceiling LEC s3
Subcapsular floor LEC
Valve LEC
UMAP
C
Macro+ medullary LEC
Ptx+ medullary LEC
Bridge LEC
Collecting vessel LEC
Oxazolone-induced LEC subset 1
Oxazolone-induced LEC subset 2
Subcapsular ceiling LEC
Subcapsular floor LEC
Transition zone LEC
Valve LEC
UMAP
B
ZMIZ1
0.8
0.7
0.6
0.5
0.4
0.3
0.2
0.1
0
UMAP
D
Zmiz1
0.6
0.5
0.4
0.3
0.2
0.1
0
UMAP

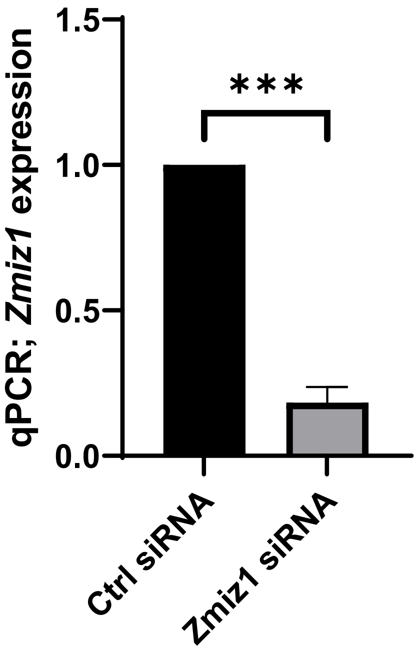

qPCR; *Zmiz1* expression
1.5
1.0
0.5
0.0
***
Ctrl siRNA
Zmiz1 siRNA

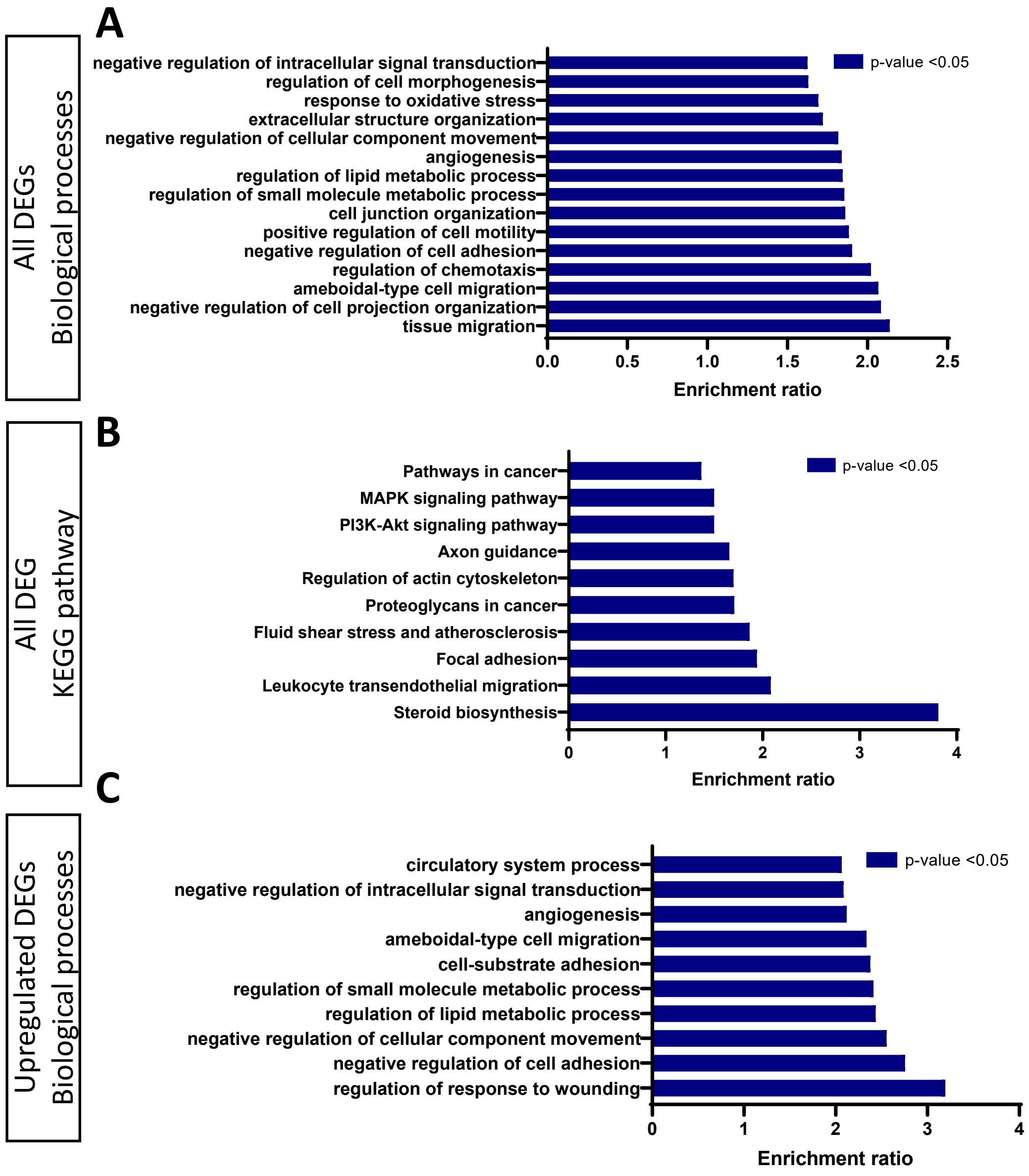
A
All DEGs
Biological processes
negative regulation of intracellular signal transduction
regulation of cell morphogenesis
response to oxidative stress
extracellular structure organization
negative regulation of cellular component movement
angiogenesis
regulation of lipid metabolic process
regulation of small molecule metabolic process
cell junction organization
positive regulation of cell motility
negative regulation of cell adhesion
regulation of chemotaxis
ameboidal-type cell migration
negative regulation of cell projection organization
tissue migration
p-value <0.05
0.0
0.5
1.0
1.5
2.0
2.5
Enrichment ratio
B
All DEG
KEGG pathway
Pathways in cancer
MAPK signaling pathway
PI3K-Akt signaling pathway
Axon guidance
Regulation of actin cytoskeleton
Proteoglycans in cancer
Fluid shear stress and atherosclerosis
Focal adhesion
Leukocyte transendothelial migration
Steroid biosynthesis
p-value <0.05
0
1
2
3
4
Enrichment ratio
C
Upregulated DEGs
Biological processes
circulatory system process
negative regulation of intracellular signal transduction
angiogenesis
ameboidal-type cell migration
cell-substrate adhesion
regulation of small molecule metabolic process
regulation of lipid metabolic process
negative regulation of cellular component movement
negative regulation of cell adhesion
regulation of response to wounding
p-value <0.05
0
1
2
3
4
Enrichment ratio

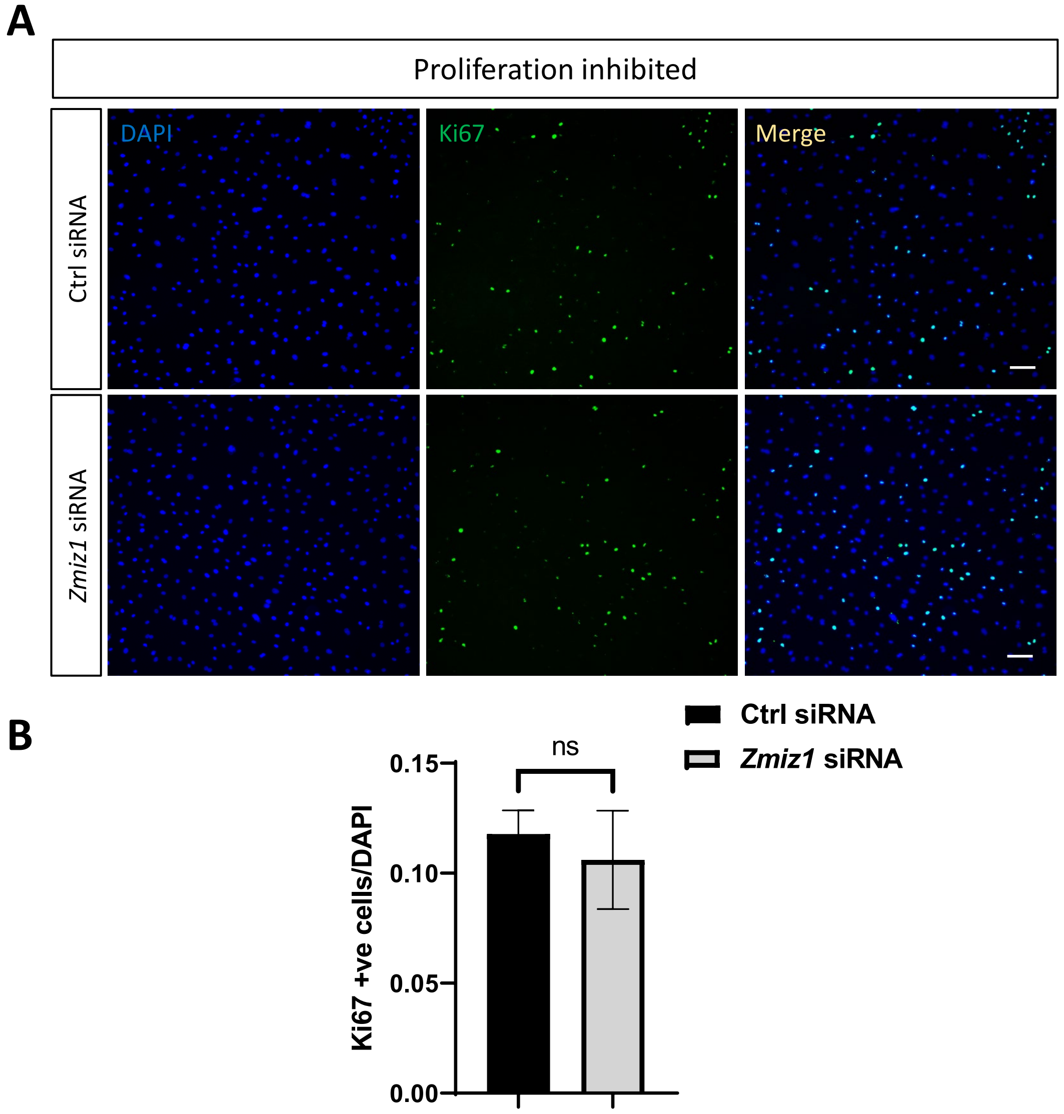
A
Proliferation inhibited
DAPI
Ki67
Merge
Ctrl siRNA
Zmiz1 siRNA
B
Ctrl siRNA
Zmiz1 siRNA
ns
Ki67 +ve cells/DAPI
0.15
0.10
0.05
0.00

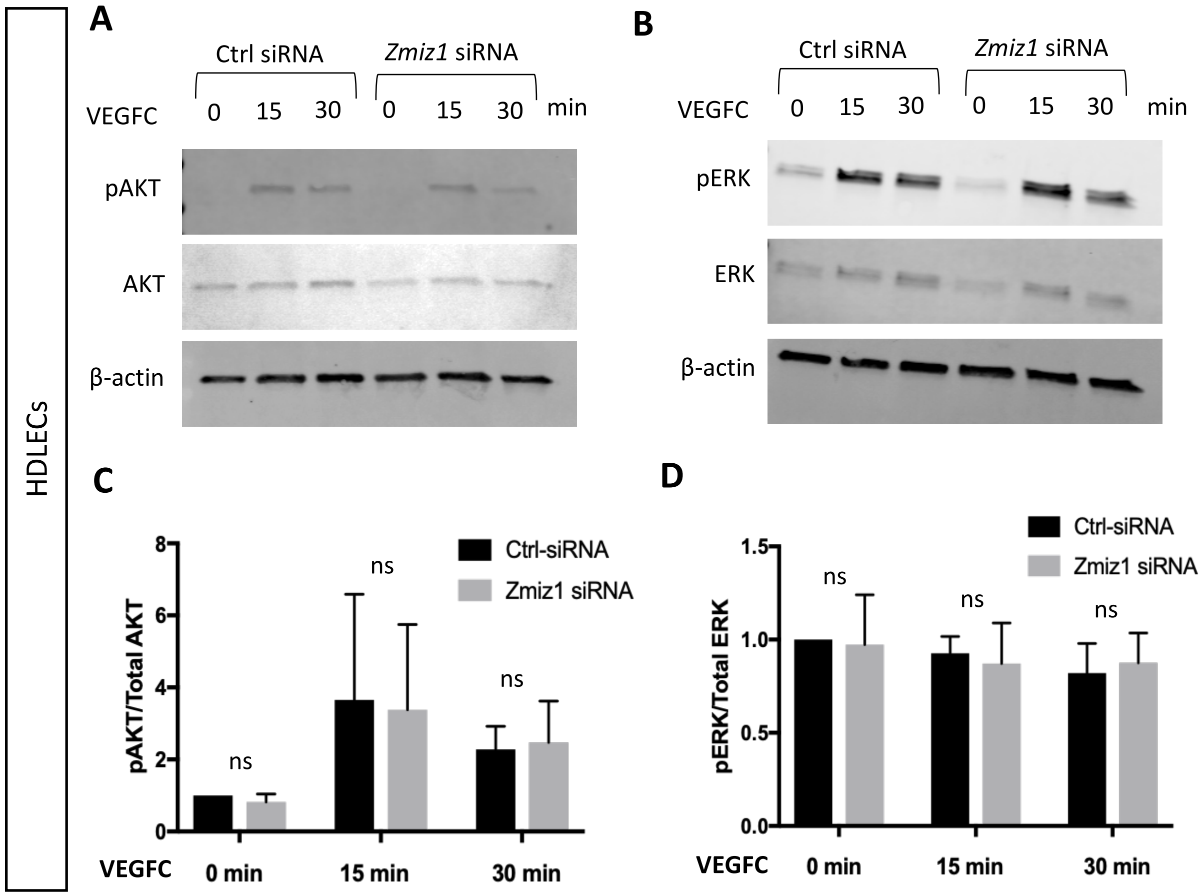
HDLECs
A
Ctrl siRNA
Zmiz1 siRNA
VEGFC 0 15 30 0 15 30 min
pAKT
AKT
β-actin
B
Ctrl siRNA
Zmiz1 siRNA
VEGFC 0 15 30 0 15 30 min
pERK
ERK
β-actin
C
Ctrl-siRNA
Zmiz1 siRNA
pAKT/Total AKT
ns
ns
ns
VEGFC 0 min 15 min 30 min
D
Ctrl-siRNA
Zmiz1 siRNA
pERK/Total ERK
ns
ns
ns
VEGFC 0 min 15 min 30 min

### ATAC seq peaks

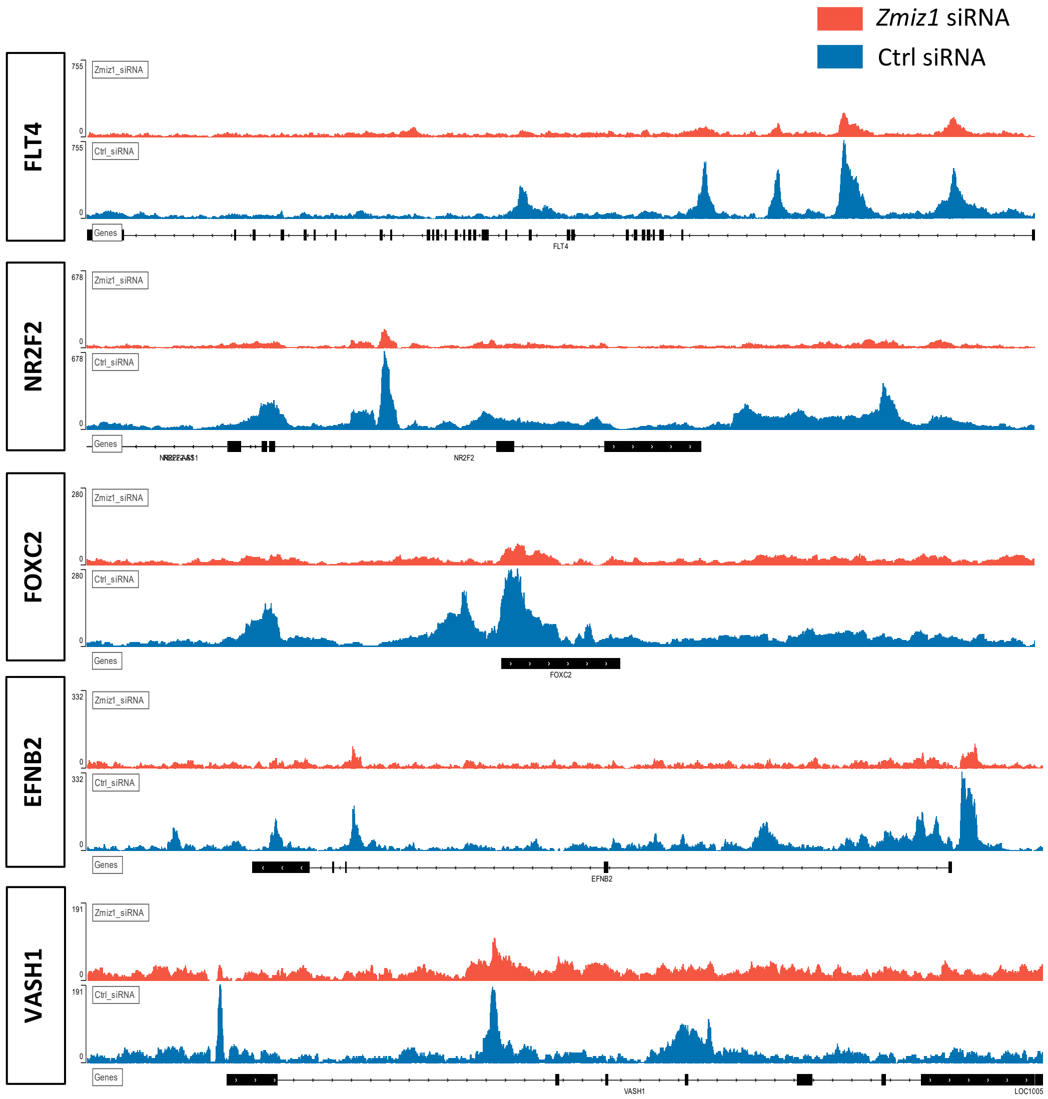

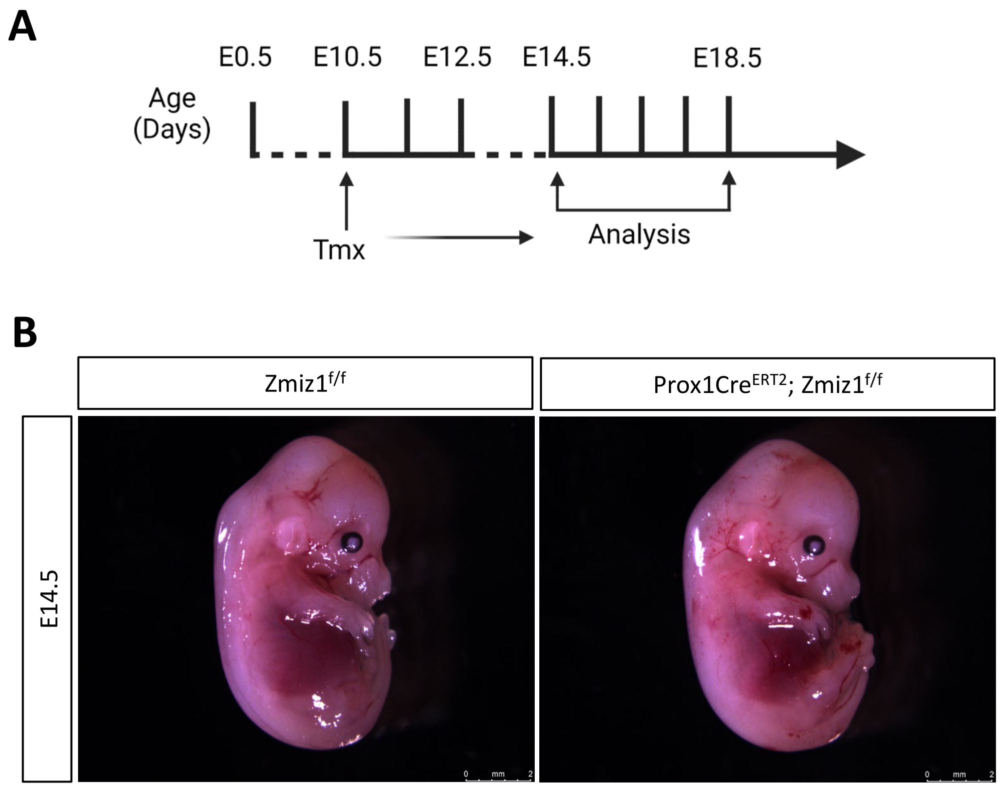

A
E0.5
E10.5
E12.5
E14.5
E18.5
Age (Days)
Tmx
Analysis
B
Zmiz1f/f
Prox1CreERT2; Zmiz1f/f
E14.5

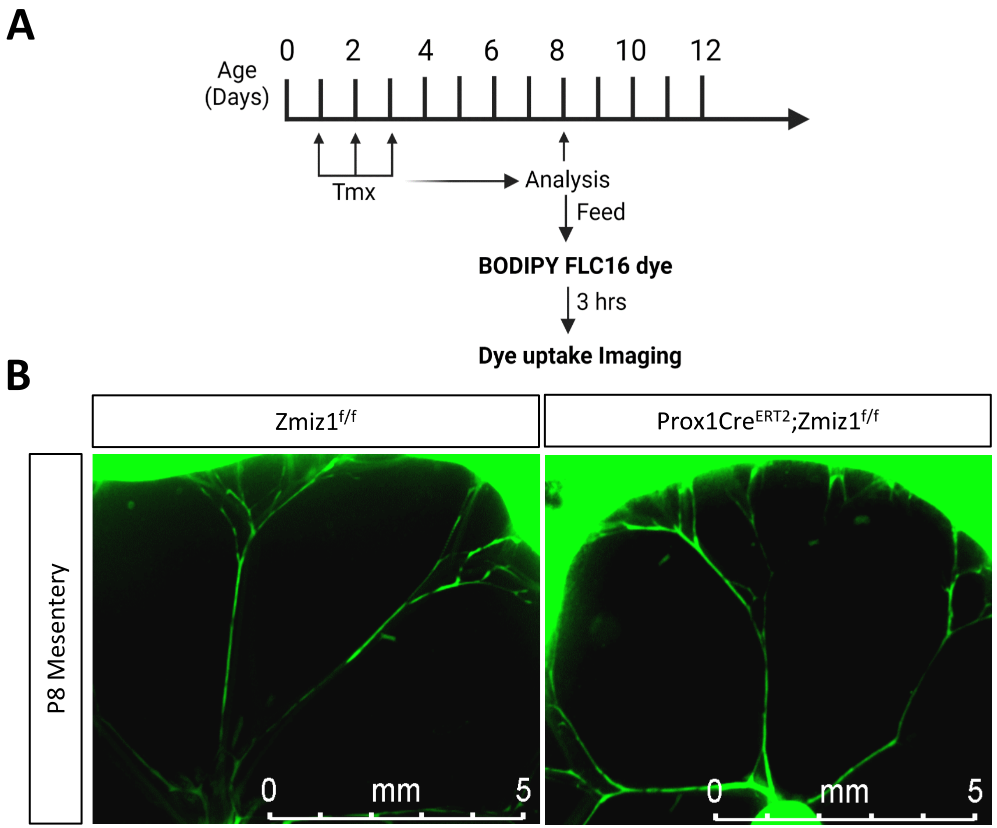
A
Age (Days)
0
2
4
6
8
10
12
Tmx
Analysis
Feed
BODIPY FLC16 dye
3 hrs
Dye uptake Imaging
B
Zmiz1f/f
Prox1CreERT2;Zmiz1f/f
P8 Mesentery
0 mm 5
0 mm 5

### Supplemental Table 1: List of qPCR primer sequence

| Gene | Forward (5’ to 3’) | Reverse (5’ to 3’) |
|---|---|---|
| PROX1 | GCTCCAATATGCTGAAGACC | CCTTGGGGATTCATGGCACTAA |
| ZMIZ1 | GTCAGCAACCATGTGTTCCACC | GCCAGTTGGTGTTCATCTGCCG |
| GAPDH | TGCACCACCAACTGCTTAGC | GGCATGGACTGTGGTCATGAG |
