## Supplemental Figure Legends for "Zmiz1 is a novel regulator of lymphatic endothelial cell gene expression and function"

**Supplementary Figure legends:**

**Supplemental figure 1**

**Zmiz1 expression comparison in mouse and human lymph node.** (A, C) Single cell UMAP of human (A) and mouse (C) lymph node lymphatic endothelial cells (LEC) subtypes. (B, D) Zmiz1 expression in distinct LEC subtypes in human (B) and mouse (D) lymph node. Adapted from https://cellxgene.cziscience.com/collections/9c8808ce-1138-4dbe-818c-171cff10e650. (Xiang et al. (2020)).

**Supplemental figure 2**

***Zmiz1* siRNA treated HDLECs exhibit depleted expression of *Zmiz1*.** (A) qPCR analysis confirms loss of *Zmiz1* expression in HDLECs treated with *Zmiz1* siRNA, as compared to control siRNA treatments (n=3).

**Supplemental figure 3**

**Gene ontology (GO) analysis of differentially expressed genes in *Zmiz1* siRNA HDLECs compared to control.** (A-B) Top biological processes (A) and KEGG pathway (B) enriched in both upregulated and downregulated genes following loss of *Zmiz1* in HDLECs. (C) Top biological processes enriched in upregulated genes following loss of *Zmiz1* in HDLECs. p-value <0.05.

**Supplemental figure 4**

**Proliferation inhibition in wound healing assay.** (A) Proliferation inhibition using Cytosine β-D-arabinofuranoside for 3 hours in HDLECs treated with ctrl and *Zmiz1* siRNA. DAPI (blue), Ki67(green). (B) Quantification for Ki67 positive (+ve) cells/DAPI (10X field) show no difference in rate of proliferation (n=3). All values mean ± SEM. Scale bars: 100 μm. ns – not significant, calculated by unpaired Student's t test.

**Supplemental figure 5**

**Unaltered response of Zmiz1 deficient LECs to VEGFC stimulation.** Western blots probed for phosphorylated(p)-AKT, AKT and β-ACTIN (A) or pERK, ERK and β-ACTIN (B) at various timepoints following VEGFC stimulations (100 ng/ml). (C) Relative quantification levels of ERK activation represented as fold change in the ratio of pAKT/total AKT relative to control treated cells at same timepoints. (D) Relative quantification levels of ERK activation represented as fold change in the ratio of pERK/total ERK relative to control treated cells at the same points. All values mean ± SEM. n = 3. ns – not significant, calculated by unpaired Student's t test.

**Supplemental figure 6**

**Reduced chromatin accessibility in lymph vessel development genes**. ATAC-seq peaks for FLT4, NR2F2, FOXC2, EFNB2, AND VASH1 in control and *Zmiz1* siRNA treated HDLECs. ATAC-seq peaks are colored blue (control (Ctrl) HDLECs) and orange (*Zmiz1* siRNA HDLECs).

**Supplemental figure 7**

**Embryonic deletion of *Zmiz1* does not lead to edema.** (A) Tamoxifen schedule used for embryonic deletion of *Zmiz1*. Tmx, tamoxifen. (B) E14.5 control and *Zmiz1*-KO embryos. No edema was observed at E14.5 and E18.5 (data not shown) (n = 3-5). Scale bars: 2 mm.

**Supplemental figure 8**

**Postnatal deletion of *Zmiz1* does not impair lymph flow.** (A) Schematic illustration of postnatal lymph flow test using BODIPY FLC16 dye. (B) Fluorescence images of P8 mesenteric lymphatic vessels in *Zmiz1*-KO pups indicate that lymph flow is not impaired after the deletion of *Zmiz1*, as BODIPY FLC16 dye was similarly present throughout the mesenteric lymphatic vessels of control mice (n = 3-5). Scale bars: 5 mm.

**Supplementary Table 1: List of qPCR primer sequence**
